# A Common Resequencing-Based Genetic Marker Dataset for Global Maize Diversity

**DOI:** 10.1101/2022.10.28.514203

**Authors:** Marcin W. Grzybowski, Ravi V. Mural, Gen Xu, Jonathan Turkus, Jinliang Yang, James C. Schnable

## Abstract

Maize (*Zea mays ssp. mays*) populations exhibit vast amounts of genetic and phenotypic diversity. As sequencing costs have declined, an increasing number of projects have sought to measure genetic differences between and within maize populations using whole genome resequencing strategies, identifying millions of segregating single-nucleotide polymorphisms (SNPs) and insertions/deletions (InDels). Unlike older genotyping strategies like microarrays and genotyping by sequencing, resequencing should, in principle, frequently identify and score common genetic variants. However, in practice, different projects frequently employ different analytical pipelines, often employ different reference genome assemblies, and consistently filter for minor allele frequency within the study population. This constrains the potential to reuse and remix data on genetic diversity generated from different projects to address new biological questions in new ways. Here we employ resequencing data from 1,276 previously published maize samples and 239 newly resequenced maize samples to generate a single unified marker set of ∼366 million segregating variants and ∼46 million high confidence variants scored across crop wild relatives, landraces as well as tropical and temperate lines from different breeding eras. We demonstrate that the new variant set provides increased power to identify known causal flowering time genes using previously published trait datasets, as well as the potential to track changes in the frequency of functionally distinct alleles across the global distribution of modern maize.

## Introduction

The degree of DNA sequence diversity observed in maize populations exceeds that of humans, most genetic model species, and many wild plants (Buckler *et al*., 2006). This diversity includes not only small scale variation – single-nucleotide polymorphisms (SNPs) and insertions/deletions (InDels) – but also copy number and presence absence variation (Swanson-Wagner *et al*., 2010). Scoring large maize populations for common sets of segregating DNA sequence polymorphisms (markers) is a key step in a range of research approaches to identify targets of selection (Hufford *et al*., 2012; Wang *et al*., 2020), inferring past demographic events, geographic diffusion (Da Fonseca *et al*., 2015; Kistler *et al*., 2018; Swarts *et al*., 2017), and linking genotype to phenotype (Mural *et al*., 2022). Early approaches to scoring common sets of genetic markers across large maize populations targeted thousands to hundreds of thousands of known markers, in the case of arrays (Ganal *et al*., 2011; Unterseer *et al*., 2014a). Array based genotyping allowed wide reuse and combination of independent datasets generated using the same array platform and, in cases where common probes were retained, between platforms. Reductions in the cost of DNA sequencing enabled sequencing-based strategies combined marker discovery and scoring in a single step (Elshire *et al*., 2011; Romay *et al*., 2013). This change reduced the substantial ascertainment bias present in many array based genetic marker datasets. However, combining marker discovery and scoring into a single step created new barriers to combining datasets. It was not possible to target specific known markers to enable interoperability between genotyping platforms. Different approaches to reducing the proportion of the genotype sequenced targeted different subsets of the genome for sequencing. Even when the same region was sequenced in two studies, differences in allele frequency, SNP calling software pipelines, or stochastic distributions of read depths might result in the same marker being identified and scored in one dataset and absent from the other. Sequencing technology has continued to improve and so costs have continued to decline. Whole genome resequencing is now economically viable for even populations of hundreds of maize genotypes. This removes the issue of generating sequence data for largely non-overlapping sites present for earlier sequencing-based strategies. However, combining marker datasets across different studies remains challenging as very different sets of markers will be discovered and pass quality filtering in different populations and/or when using different bioinformatics pipelines.

Identifying a common set of genetic variants in maize is challenging, and the optimal set of lines to use in defining a marker set likely depends on the question of interest. Maize was domesticated from a wild progenitor teosinte (*Zea mays* ssp. *parviglumis*) 9,000-10,000 years ago in southwest Mexico (Matsuoka *et al*., 2002; Piperno *et al*., 2009) with substantial gene flow from at least one other teosinte (*Zea mays* ssp. *mexicana*) (Chen *et al*., 2022; Van Heerwaarden *et al*., 2011). After domestication, maize spread across North and South America (Da Fonseca *et al*., 2015; Kistler *et al*., 2018; Swarts *et al*., 2017). Maize, almost certainly of Caribbean origin, was first cultivated in southern Europe in 1493 and was growing in Germany by 1539 (Tenaillon and Charcosset, 2011). By 1555, substantial maize cultivation was already being recorded in Henan, China (Ho, 1955). Therefore, maize was already cultivated on at least four continents in the mid XVI-century. Tropical maize varieties that flower under short-day conditions making them unsuitable for cultivation in regions with killing frosts retain many alleles and haplotypes not found in temperate populations (Hung *et al*., 2012). Breeding efforts in the United State, Europe, and China focus on temperate-adapted cultivars which are less photoperiod sensitive than tropical maize. In the United States, hybrid production focuses on three heterotic groups, stiff stalk, non-stiff stalk, and iodent, in Europe many hybrids are generated from crosses between the flint and dent heterotic groups, while in China Huangzaosi group was also used alongside stiff stalk, non-stiff stalk, and iodent (Wang *et al*., 2020). As a result, different research groups studying quantitative genetic variation, domestication, adaptation, or crop improvement have selected different sets of inbred lines, open-pollinated landraces, or maize wild relatives drawn from populations in different parts of the globe.

The maize HapMap2 project was motivated in part by understanding changes in genetic diversity associated with maize domestication and improvement. The study identified more than 55 million total variants from an average of 4x resequencing of 103 samples, including 83 individuals representing domesticated maize and 20 individuals drawn from wild relative populations aligned to B73_RefGen_V1 (Chia *et al*., 2012; Hufford *et al*., 2012). A project focused on understanding the history and demography of the initial introduction of maize to Europe identified 22.3 million SNPs relative to the B73_RefGen_V2 genome by resequencing 67 maize samples originating in the Americas (n=37) and Europe (n=30) to an average depth of 18x (Brandenburg *et al*., 2017). Given the focus on the introduction of maize to Europe, this study focused primarily on maize lines originating in western (18) and central (11) Europe, with one line sourced from eastern Europe. Another study focused on the pre-Colombian demographic history of maize resequenced 35 maize landraces and wild relatives from the Americas to a median depth of 28x and identified 49.5 million SNPs via alignment to the B73_RefGen_V3 reference genome (Wang *et al*., 2017). A study of maize domestication and improvement in South America generated data from 49 living and archaeological maize samples and generated a new SNP set by aligning both data from these new samples and 70 published maize datasets to the B73_RefGen_V4 reference genome (Kistler *et al*., 2018). Resequencing of 521 entry maize association panel to an average depth of 20x identified 11.5 million variants as part of an effort to link structural variation in the genome to changes in gene expression and phenotypic outcomes (Yang *et al*., 2019). A comparative analysis of phenotypic and genetic changes associated with the breeding effort in different temperate breeding programs generated resequencing data from 350 maize inbreds from China (187) and the United States (163) sequenced to a median depth of 12x and identified more than 29 million genetic markers relative to the B73_RefGen_V3 reference genome (Wang *et al*., 2020). An effort to quantify SNP and transposon insertion diversity within an association panel used for genome wide association studies identified approximately 2.4 million SNPs and 0.45 million segregating transposon associations across a panel of approximately 500 temperate adapted maize lines (Qiu *et al*., 2021; Renk *et al*., 2021). Finally, a recent study of genus wide genetic variation in maize identified approximately 65 million SNPs and approximately 8 million InDels by generating 22x average depth sequencing data from 239 accessions of wild relatives (Chen *et al*., 2022; Gui *et al*., 2022) in combination with the approximately 500 entry maize diversity panel resequenced in (Yang *et al*., 2019). The largest scale of these efforts to date is likely the aggregate analysis of 1,218 maize lines as part of the maize HapMap3 project representing global maize diversity, however, higher sequencing costs at the time of this study resulted in lines being resequenced to a median depth of 2x (Bukowski *et al*., 2018).

Here we sought to update and expand the reference set of segregating diversity in maize by incorporating published high coverage resequencing data from maize lines originating on six continents, including resequencing data from lines relative to maize domestication and improvement, including wild relatives, tropical landraces, and archaeological maize samples, as well as maize wild relatives, and to further improve the resolution and mapping power for maize genome wide association studies conducted in the temperate midwest through the resequencing of an additional 239 maize lines including 228 lines from the Wisconsin Diversity panel not previously resequenced and 11 Eastern European lines. To ensure the greatest degree of reusability and forward compatibility, we employed the B73_RefGen_V5 maize reference genome (Hufford *et al*., 2021) and, in addition to raw and filtered SNP files, we are releasing GATK GenomicsDB datastores so that these same 1,515 lines can be incorporated into future high coverage maize resequencing efforts without the need to reprocess and realign sequence data.

## Methods

### Plant material and datasets

Whole genome resequencing data from 1,515 total samples were used in this analysis, including 1,276 previously published samples (Brandenburg *et al*., 2017; Bukowski *et al*., 2018; Chen *et al*., 2022; Chia *et al*., 2012; Kistler *et al*., 2018; Qiu *et al*., 2021; Unterseer *et al*., 2014b; Wang *et al*., 2020, 2017) and 239 lines resequenced as part of this study. The origin and source of each sample included in this analysis are provided in supplemental table S1.

Two hundred twenty-eight inbred lines from the Wisconsin Diversity Panel (Mazaheri *et al*., 2019) were grown in a greenhouse setting (27°C - 29°C during the day, and 19°C – 21°C at night, with 12 hours light/12 hours dark). After reaching V2, the youngest leaf was harvested onto the ice and was then lyophilized for two days in a Flexi-Dry lyophilizer (FTS Systems Inc. New York). The lyophilized samples were ground to a fine powder at room temperature using 4.76 mm ball bearings in a Tissuelyzer II (Qiagen, Germany). Following the manual’s instructions, DNA was extracted from the individual lyophilized and ground samples using the MagMAX Plant DNA Isolation Kit (Thermo Scientific, USA) with the help of a benchtop automated extraction instrument, KingFisher Flex (Thermo Scientific, USA). Raw DNA extracts were quantified using the Quant-iT dsDNA Broad Range Kit (Invitrogen, USA) and normalized to 20ng/uL using an Andrew pipetting robot (Andrew Alliance, USA). Normalized DNA samples were submitted to Psomagen, Inc. (USA) for library preparation and sequencing.

On receipt of the DNA samples, Psomagen, Inc. (USA) performed an in-house Quality assessment of the DNA samples using TapeStation 4200 (Agilent). DNA samples from 29 lines did not meet the minimum DNA quality control standards for sequencing. An additional set of seeds from these lines were surface sterilized by washing them in a 5% v/v bleach solution for 10 minutes, rinse three times with sterile water, and placed in centrifuge tubes with wetted paper and left in the dark at 23°C. Shortly after germinating (VE), the entire coleoptile was harvested, snap-frozen in liquid nitrogen, and stored at -80°C. The tissue was then ground to a fine powder using 3/16 inch (4.76mm) ball bearings in a Tissuelyzer II (Qiagen, Germany) in the presence of dry ice in the pockets around tube holders. The DNA extraction was then performed utilizing the same procedure as was used on the original samples. Initial quality assessment of DNA samples, library preparation, and sequencing was performed by Psomagen, Inc. (USA). Libraries were prepared using the TruSeq DNA PCR-Free kit (Illumina, USA). A NovaSeq6000 S4 (Illumina, USA) sequencer was used to generate 150 bp paired-end reads.

Eleven Polish inbred lines were obtained from Plant Breeding Smolice Ltd., Co., Poland. Plants were grown in a phytotron chamber (24°C/22°C day/night and 16 hours light/8 hours dark). Tissues for DNA extraction were harvested from the third fully developed leaf (V3 stage) and three individual plants were pooled into a single sample. Leaves were immediately flash-frozen in liquid nitrogen and tissue was ground in liquid nitrogen using a mortar and pestle. DNA extraction was done with DNeasy Plant Kit (Qiagen, Germany) according to the manual’s instructions. Genomic DNA for each genotype was submitted to Fasteris (Switzerland) for whole genome sequencing. For S160, S50676, and S68911 inbred lines 100 bp paired-end reads, and for the remaining eight lines 150 bp paired-end reads were generated on a HiSeq X Ten (Illumina, USA) sequencer.

### Creation of the global maize SNP set

After fastq files were downloaded from the European Nucleotide Archive or transferred from the sequencing provider, each file was cleaned using fastp v.0.23.2 with the default setting (Chen *et al*., 2018). Reads with > 40% unqualified bases or quality value < 15 were removed. Cleaned fastq files were aligned to the B73_RefGen_V5 maize reference genome (Hufford *et al*., 2021) using SpeedSeq v.0.1.2 (Chiang *et al*., 2015) which parallelizes BWA-MEM v.0.7.10 (Li, 2013) for alignment, Samblaster v.0.1.22 for marking duplicated reads (Faust and Hall, 2014), and Sambamba v.0.5.9 for position sorting and BAM file indexing (Tarasov *et al*., 2015). Samblaster defined duplicate read pairs as cases where two or more pairs of reads aligned to the same reference sequence on the same strand and with the same 5^′^ start position – or inferred 5^′^ start position if the alignment was clipped – for both forward reads and for both reverse reads. Unless otherwise stated, default parameters were used for each software package.

Individual gVCF files were generated for each maize pseudomolecule for each BAM file using the Haplotype-Caller tool provided by GATK v.4.2.0.0 in diploid mode (Poplin *et al*., 2018). To enable extensive parallelization of variant calling the maize genome was divided into 5 Mb windows for the creation of separate GenomicsDB datastores. During the project, an update of GATK appeared (v.4.2.6.1), which offered a reduction of files number stored in GenomicsDB datastores. Therefore, GenomicsDBImport tool provided by GATK v.4.2.6.1. was used for each genomic window to create GenomicsDB datastore. Joint variant calling was conducted using the GenotypeGVCFs tool provided by GATK v.4.2.6.1. with default settings. To aid in additional parallelization, each 5 Mb GenomicsDB datastore was divided into five 1 Mb windows for variant calling.

Following GATK best practices recommendations, hard filters were applied to call variants. Variants were divided into SNPs and InDels for filtering. SNPs having a QualByDepth < 2.0, FisherStrand > 60.0, RMSMappingQuality < 40.0, MappingQualityRankSumTest < -12.5 or ReadPosRankSumTest < -8.0 were removed. InDels with QualByDepth < 2.0, FisherStrand > 200.0 or ReadPosRankSumTest < -20.0 were also removed. After filtering, SNP and InDel variants were merged into single sorted VCF files for each chromosome using Picard v.2.9 (Pic, 2019). Finally, genotypes with depth < 2 were masked using bcftools setGT plugin v.1.10.2 (Danecek *et al*., 2021). All further VCF files manipulations were done with bcftools v.1.10.2 (Danecek *et al*., 2021).

### Creation of the filtered and imputed maize SNP sets

The filtered and imputed variant set was generated by first removing variants where: >2 alleles were observed in the population, variants with ≥ 50% missing data, variants with extremely low < 1,515 or extremely high > 33,550 sequencing depth, and variants with inbreeding coefficients ≥ 0 resulting in ∼ 46 million variants. The inbreeding coefficient per variant was calculated as:

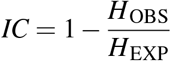

where H_obs_ and H_exp_ are the observed and expected heterozygosity under Hardy–Weinberg equilibrium.

Variants were phased and imputed using Beagle 5.0 with default settings (err=0.0001; window=40.0 cM; overlap=4.0 cM; step=0.1 cM; nsteps=7) (Browning *et al*., 2018).

### Population genetic analyses

Principle component analysis (PCA) was conducted with Plink v.1.9 (Purcell *et al*., 2007). Unimputed variants were filtered with MAF > 5% and with a fraction of missing data < 10%, leading to 19,205,674 markers, which were used for PCA. Individual genotypes were assigned to the population using published literature data (Brandenburg *et al*., 2017; Bukowski *et al*., 2018; Chen *et al*., 2022; Chia *et al*., 2012; Kistler *et al*., 2018; Qiu *et al*., 2021; Unterseer *et al*., 2014b; Wang *et al*., 2020, 2017).

Measures of LD (*r*^2^) were calculated for the entire population and pre-defined groups using PopLDdecay v.3.42 (Zhang *et al*., 2019), and the subset of unimputed SNPs with MAF > 0.05 and missing rate < 0.25. Local LD in a 100 Kb window was calculated using the Genome-wide Complex Trait Analysis (GCTA) with the default settings (Yang *et al*., 2011).

To calculate nucleotide diversity (Nei and Li, 1979) maize genome was first divided into 1 Kb window using bedtools v.2.27.1 (Quinlan and Hall, 2010). Next, windows that overlapped with region annotated as transposon in B73_RefGen_V5 (Hufford *et al*., 2021) were excluded from the analysis. Nucleotide diversity were calculated using the vcftools v.0.1.16 site-pi function (Danecek *et al*., 2011) in remaining 1 Kb window, with a value of 0 employed for monomorphic positions. Mean values were calculated for each window and the distribution of window-mean values were employed in downstream analyses.

### Genome-wide association study

A published dataset of female flowering time (days to silking) for 752 inbreds drawn from the Wisconsin Diversity panel (Mazaheri *et al*., 2019) and grown in a replicated field study in Lincoln, NE in 2020 was employed for genome wide association (Mural *et al*., 2022). Three genetic marker sets for the same population of 752 maize inbreds were used to conduct GWAS. The first set was created by filtering 752 maize inbreds with MAF > 5% from 899,784 variants called using RNA-seq and called relative to the B73_RefGen_V4 reference genome (Mazaheri *et al*., 2019). This leads to a creation set containing 428,487 variants. The second was a set of ∼17.2 million markers called using a combination of resequencing (581 lines) (Bukowski *et al*., 2018; Qiu *et al*., 2021) and RNA-seq (361 lines) (Hirsch *et al*., 2014; Mazaheri *et al*., 2019) with extensive imputation to fill in non-exonic SNPs for the subset of samples genotyped only with RNA-seq (Mural *et al*., 2022; Sun *et al*., 2022). The third was a set of 16,634,049 markers obtained by subsetting the filtered and imputed SNP set assembled in this study to include only those markers with a minor allele frequency >5% among the 752 genotypes for which female flowering time phenotypes were available. In all three cases, GWAS was conducted using the mixed linear model algorithm (Yu *et al*., 2006) as implemented in the rMVP R package v.1.0.6 (Yin *et al*., 2021). In all three cases, both the kinship matrix – computed following the method described in (VanRaden, 2008) – and the first five principal components of variation – calculated as described above – were included as covariates. Calculations of local linkage disequilibrium were performed using Plink v.1.9 (Purcell *et al*., 2007).

All additional statistical analysis were conducted in R (R Core Team, 2022), with extensive use of *data*.*table* (Dowle and Srinivasan, 2021), and *tidyverse* (Wickham *et al*., 2019) for data manipulation, and *tidyverse* and *patchwork* (Pedersen, 2020) for visualisation.

## Results and discussion

### Sequence Variation Across the Genome of Maize

Sequence data from 1,276 maize individuals generated as part of eight different studies (Brandenburg *et al*., 2017; Bukowski *et al*., 2018; Chen *et al*., 2022; Chia *et al*., 2012; Kistler *et al*., 2018; Qiu *et al*., 2021; Unterseer *et al*., 2014b; Wang *et al*., 2020, 2017) were retrieved from the European Nucleotide Archive. To this public dataset, we added data from *de novo* resequencing of 228 maize inbred lines which are part of the expanded Wisconsin Diversity Panel (Mazaheri *et al*., 2019) but were not resequenced as part of previous efforts (Bukowski *et al*., 2018; Qiu *et al*., 2021). An average of 155 million reads were generated for each of these inbred lines, corresponding to an average sequencing depth of approximately 22x. Additionally, a set of 11 maize inbred lines from Poland, representative of eastern Europe, a region only modestly represented among previous maize resequencing efforts were resequenced here to an average depth of ∼35x. Those lines were used in previous studies on maize cold response (Grzybowski *et al*., 2019; Sowiński *et al*., 2005). The total set of 1,515 maize accessions included wild relatives, archaeological samples, modern open-pollinated varieties, and inbred lines from both public and private sector breeding efforts representing the maize lines originating in or developed over six continents (Table S1).

Aligning sequence data from each of these accessions to the maize B73_RefGen_5 reference genome (Hufford *et al*., 2021) and applying recommended filtering criteria from GATK resulted in the identification of 365,611,965 potential DNA sequence polymorphisms. This number is substantially higher than ∼83 million variants identified in the maize HapMap3 project, one of the largest surveys of maize genetic diversity conducted to date, incorporating data from 1,218 maize accessions (Bukowski *et al*., 2018). However, it should be emphasized that HapMap3 utilized a different variant calling pipeline and that the median sequencing depth of samples in that study was ∼2x. A more recent study that examined the genetic differentiation of male and female heterotic groups in maize using resequencing data from 1,604 maize inbred lines, primarily from China and the USA, resequenced to an average depth of ∼7.5x identified ∼242 million DNA sequence polymorphisms (Li *et al*., 2022).

Second-stage quality filtering (based on allele number, missing data rates, sequence depth, and excess heterozygosity, see Methods) resulted in a smaller set of 46,054,265 higher confidence variants, including 43,296,332 SNPs and 2,757,933 InDels (Figure. S1). The median total sequencing depth for higher confidence variants was 17,365 (Figure. S2), corresponding to an average sequence depth of 11.5 reads per site per individual. Concordance rates for SNP calls among the 26 NAM founder parents (Hufford *et al*., 2021) and SNP calls reported as part of the *de novo* sequence assembly of these parents ranged from 92% to 99% with a mean value of 98% (Table S2). Among these higher confidence variable sites, the median accession was genotyped as heterozygous 2.8% of the time. However, per-accession heterozygosity rates varied significantly across groups (Figure S4). Heterozygous calls were more common in pericentromeric regions (Figure S4). Groups expected to consist primarily of inbred lines, such as those classified as belonging to the stiff stalk, non stiff stalk, and iodent heterozygous groups typically exhibited per-accession heterozygosity values of <3%. Accessions classified as wild-relatives frequently exhibited per-accession heterozygosity values of >10% (Figure S4, Table S1). Inbred lines with unexpectedly high heterozygosity were not removed from the final dataset however they should be used with caution as these may represent contaminated or mislabeled samples.

While many high confidence SNPs (41%) and InDels (38%) were rare, defined here as a minor allele frequency ≤ 5%, more than 26 million variants were common defined as a minor allele frequency >5% (25,154,632 SNPs and 1,704,190 InDels) (Figure 1b&d). Segregating SNPs were more common around pericentromeric regions (Figure 1a) while segregating InDels were more frequent on chromosome arms and less frequent in pericentromeric regions (Figure 2c). The relationship between distance from the centromere and SNP or InDel density was extremely weak but statistically significant for each chromosome (Figure S5), similar to the pattern of SNPs and InDels reported in sorghum (Lozano *et al*., 2021). Linkage disequilibrium was typically elevated in pericentromeric regions likely reflecting lower recombination rates in these regions (Figure 1e). The pattern of elevated linkage disequilibrium around the centromere was less prominent on chromosome 10, consistent with previous reports (Romero Navarro *et al*., 2017). Several other peaks of elevated linkage disequilibrium were observed which did not coincide with the known positions of maize centromeres. One potential explanation is that these peaks may represent large segregating structural variants (Crow *et al*., 2020) however validating hypothesis is beyond the scope of this paper. The majority of high confidence variants (57%) were located in intragenic regions, defined as those regions ≥ 5 Kb from the closest annotated exon. Another 31% of variants were located in regions outside annotated genes but < 5 Kb from the closest gene (Figure 1f). Among variants located between the annotated transcription start sites and transcription stop sites of genes, intronic variants were most abundant (8.6%), followed by 5^′^- and 3^′^ UTR (0.7 and 0.9%) and coding sequence (0.4%).

**Figure 1.**
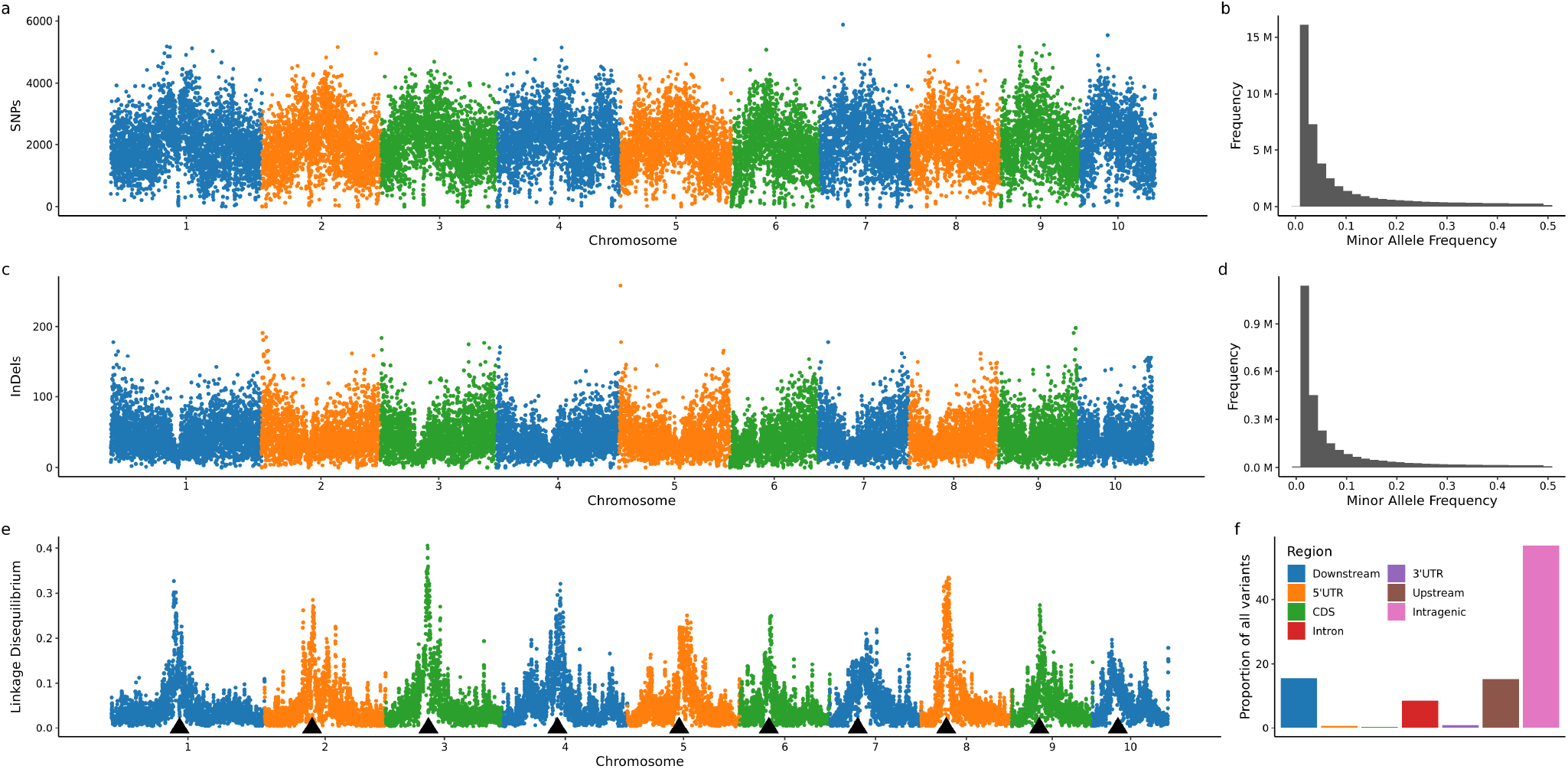
Properties of high confidence maize genetic variants identified in this study. The distribution of high confidence, and common (MAF > 5%, ∼27 million) **(a)** SNPs and **(c)** InDels across each of the 10 maize chromosomes. For both **(a)** & **(b)** the genome was divided into non-overlapping 100 Kb windows and SNPs and InDels were counted in each window. Distribution of minor allele frequency of high confidence (∼46 million) **(b)** SNPs and **(d)** InDels. **(e)** Mean LD value in 100 Kb window calculated with high confidence, and common (MAF > 5%) SNPs. Black triangles indicate the centromere position on each chromosome. **(f)** Percentage of variants across the major genic and intergenic regions calculated with high confidence variant set.

**Figure 2.**
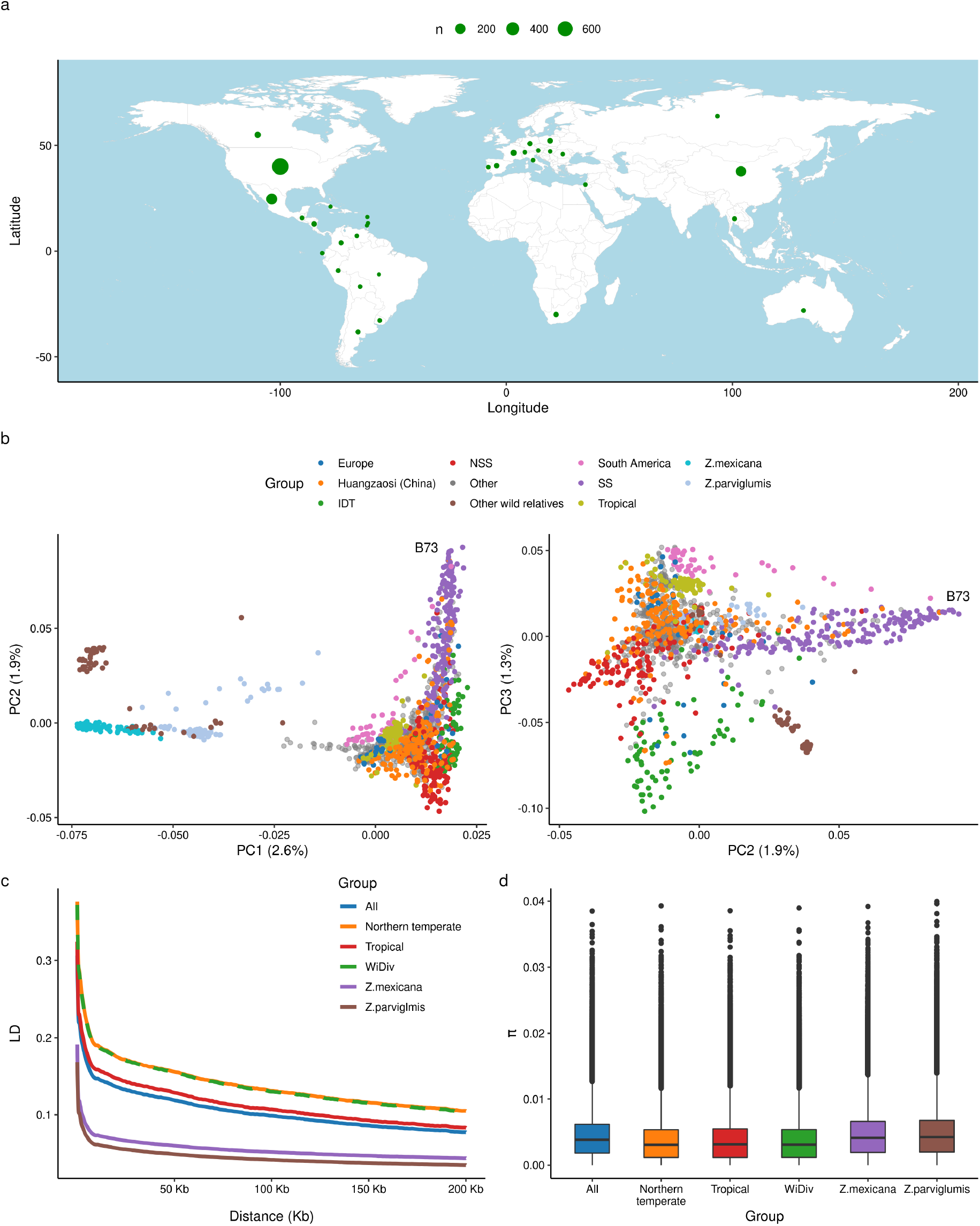
Geographical distribution, population structure, LD patterns, and nucleotide diversity in maize. **a** Geographical distribution of the country of origin for 1,515 maize individuals.**b** First three principal components from PCA analysis on 1515 maize individuals. Each individual was assigned to different groups based on previous literature data. **c** Genome-wide averaged distance of LD decay for six maize groups. **d** Nucleotide diversity for six maize groups. High confident common (MAF > 5%) variant set were used for each analysis.

### Intra- and Inter-Population Genetic Variation

Of the 1,515 maize samples used in this study, 760 were assigned to one of ten groups through a combination of prior publication data and metadata associated with USDA GRIN records. These ten groups included three groups of wild relatives – *Zea mays ssp. mexicana* (n=79, hereafter *mexicana*), *Zea mays ssp. parviglumis* (n=84, hereafter *parviglumis*), and other wild relatives (n=66). Among these samples, four groups based on geographic origin – tropical (n=86), South America (n=48), China (n=182), Europe (n=34) – and three based on a combination of geographic origin and heterotic group – temperate North American stiff stalk (n=193), non-stiff stalk (n=127), and iodent (n=69). The remaining 549 lines were classified as “other” in the analyses below. All together, this set of lines comes from 35 countries across six continents (Figure 2a). Lines tended to cluster based on the group assignment in analyses of population structure conducted using the genetic marker data generated in this study (Figure 2b and S6). The first principal component of variation for genetic marker data roughly corresponded to the division between the maize and other maize wild relatives (Figure 2b). The second principal component separates stiff stalk and non-stiff stalk heterotic groups, alternatively, to how closely a line is related to B73, the reference genotype for maize. Finally, the third principal component corresponds to latitudinal geographic distribution, with South American lines at one extreme, followed by tropical and wild populations, then Chinese, European, and North American temperate populations, and other wild relatives (Figure 2b).

High-density genetic marker data is useful for both population genetic and quantitative genetic analyses (Mural *et al*., 2021). Many population genetic analyses require measurements of plant traits. When trait data is collected in different environments, variance resulting from differences in genotype is confounded with variance resulting from different environments, reducing statistical power to link genotype and phenotype. Growing and phenotyping large plant populations in common environments can more effectively isolate contributions of genetic variation to phenotypic variation, at least in that specific environment. However, this presents a challenge in capturing global genetic diversity in species such as maize where different lines are adapted to different environments and may not even be able to successfully complete their lifecycles in environments to which they are not adapted. Efforts to establish common association panels for quantitative genetic analysis in maize including the Maize Association Panel (MAP) (Flint-Garcia *et al*., 2005), Shoot Apical Meristem association panel (SAM) (Leiboff *et al*., 2015), and the Wisconsin Diversity Panel (WiDiv) (Hansey *et al*., 2011) have required researchers to prioritize the partially contradictory goals of maximizing genetic diversity while also selecting for a set of genotypes that can all grow and successfully complete their life cycles in a single common environment. Based on marker data for 798 genotypes from the WiDiv panel included in this study, linkage disequilibrium decays roughly as fast within the WiDiv panel as with the set of all northern temperate lines (1090 lines defined as all those excluding teosinte, tropical, and South America lines) but mostly more slowly than the rate of linkage disequilibrium decay among all 1,515 lines included in this study (Figure 2c). LD decayed fastest among the two maize wild relative populations with the largest number of samples: *mexicana* and *parviglumis*. The median value of *π* observed for randomly selected intervals in the maize genome within the WiDiv population was 0.00376, similar to the median observed for all temperate lines (*π*=0.00374), but lower than when calculated for the population of all genotypes included in this study (*π*=0.00443; Figure 2d). The difference in *π* for the overall population is likely driven by the inclusion of wild relatives in the overall population as these populations exhibit elevated *π* values of *mexicana* (*π*=0.00480) and *parviglumis* (*π*=0.00490). Previous study indicate that 83% of nucleotide variation from *teosinte* being retained in maize landraces (Hufford *et al*., 2012). Here we found that WiDiv lines retain 76% of nucleotide variation observed in *parviglumis*, which indicate that a substantial portion of genetic variation is still present in this northern temperate set of lines.

### Greater Utility of Existing Trait Data as Marker Density Increases

Community association panels are typically reused by many research groups working to study the genetic control of variation in different traits of interest. A recent literature study identified more than 160 distinct trait datasets scored across North American temperate maize association panels between 2010 and 2020 (Mural *et al*., 2022). During the past seventeen years, the density of publicly available markers for maize association panels has grown from 94 microsatellite markers (Flint-Garcia *et al*., 2005) to 1,536 microarray-based SNP markers (Hansey *et al*., 2011) to hundreds of thousands of markers scored using genotyping by sequencing (Romay *et al*., 2013) and approximately one million markers scored using RNA-seq (Leiboff *et al*., 2015; Mazaheri *et al*., 2019) and now typically include tens of millions of markers discovered and scored via whole genome resequencing (Bukowski *et al*., 2018; Chen *et al*., 2022; Li *et al*., 2022; Qiu *et al*., 2021; Wang *et al*., 2020) or a combination of whole genome resequencing for a subset of lines and imputation from lower density markers for additional lines (Mural *et al*., 2022; Sun *et al*., 2022).

We employed a previously published set of female flowering data (days to silking) generated for 752 temperate adapted maize inbreds to assess the impact of increased marker density vs direct resequencing (this study) on the outcomes from genome wide association studies in maize. When using ∼400k markers discovered and scored using RNA-seq (MAF > 5% in 752 lines) (Mazaheri *et al*., 2019), a genome wide association study identified one statistically significant signal corresponding to the cloned maize flowering time gene *MADS69* ((Liang *et al*., 2019); Figure 3a). A genome wide association study conducted using the new, purely whole genome resequencing based marker dataset generated in this study identified both *MADS69* and *ZCN8* (Figure 3b). Overall, the newly generated variant dataset increases the power to detect causal genes and doesn’t increase p-value inflation (Figure S7).

**Figure 3.**
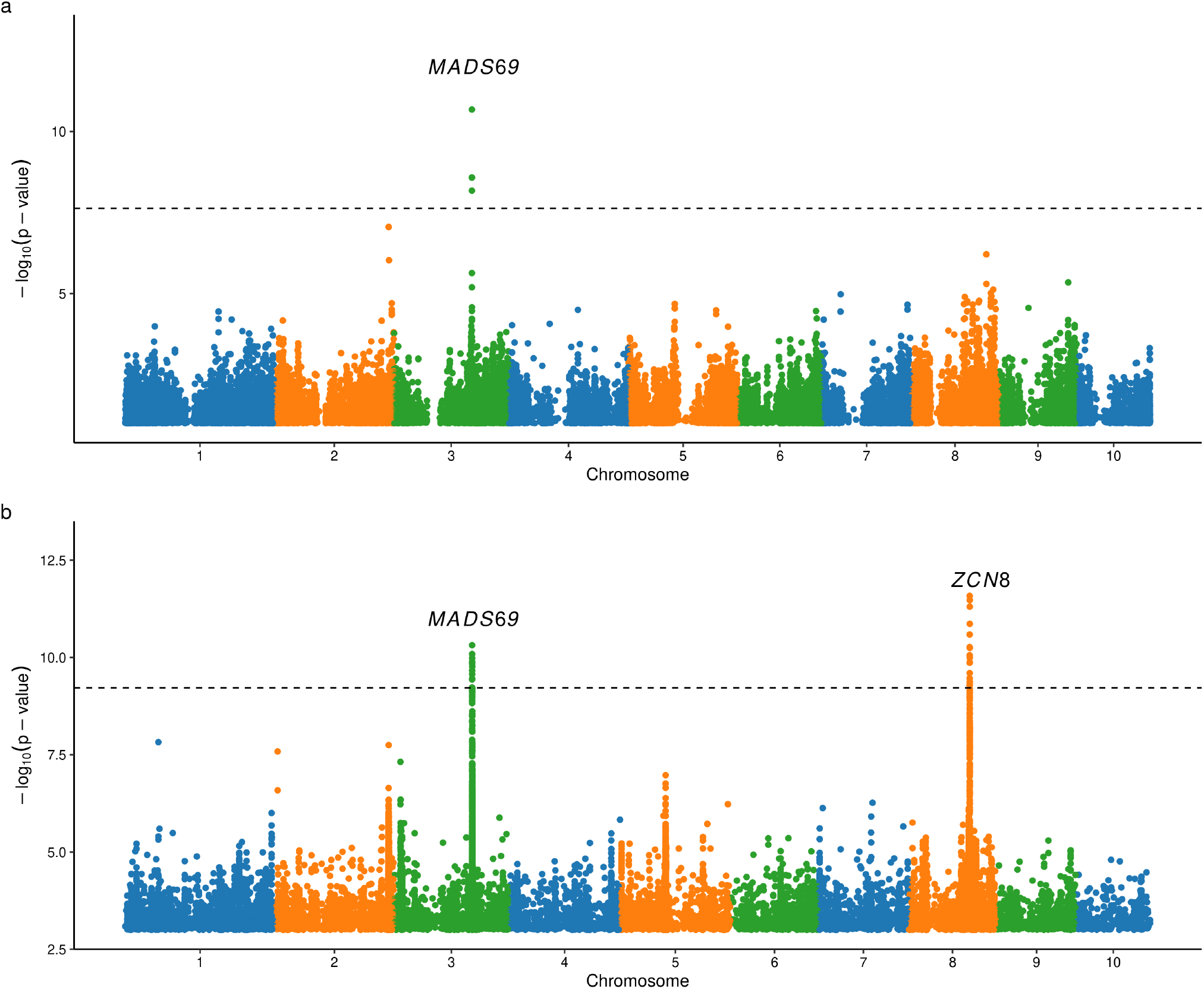
Identification of the candidate genes for flowering time (days to silking) via GWAS. **a** Association between days to silking, as reported in Mural *et al*. (2022), and 428,487 segregating SNPs identified and genotyped using RNA-seq data in Mazaheri *et al*. (2019).**b** Association test for days to silking using the marker set defined in this study (n = 16,634,049). The horizontal dashed line on each plot indicates an *α* = 1% significance threshold after applying Bonferroni correction assuming n number of variants in each dataset as independent tests.

In addition to the total number of confident signals identified, an additional potential benefit of higher-density genetic marker data is the more precise localization of peaks to only one or several candidate genes. The peak corresponding to *MADS69* included 29 markers which were significant at a Bonferroni corrected p-value of 0.01. These markers span a region of 410,350 bp that includes 3 annotated genes. However, the peak SNP (e.g. the single SNP with the most significant p-value) was 8,507 bases from *MADS69* and *MADS69* was the closest gene to this SNP (Figure 4a). The peak corresponding to *ZCN8* included 35 significant markers which were significant at a Bonferroni corrected p-value of 0.01. These markers span a region of 349,944 bases that includes 7 annotated genes. In this case the peak SNP was 18,912 bases from *ZCN8* and three genes separated *ZCN8* from the peak SNP (Figure 4b).

**Figure 4.**
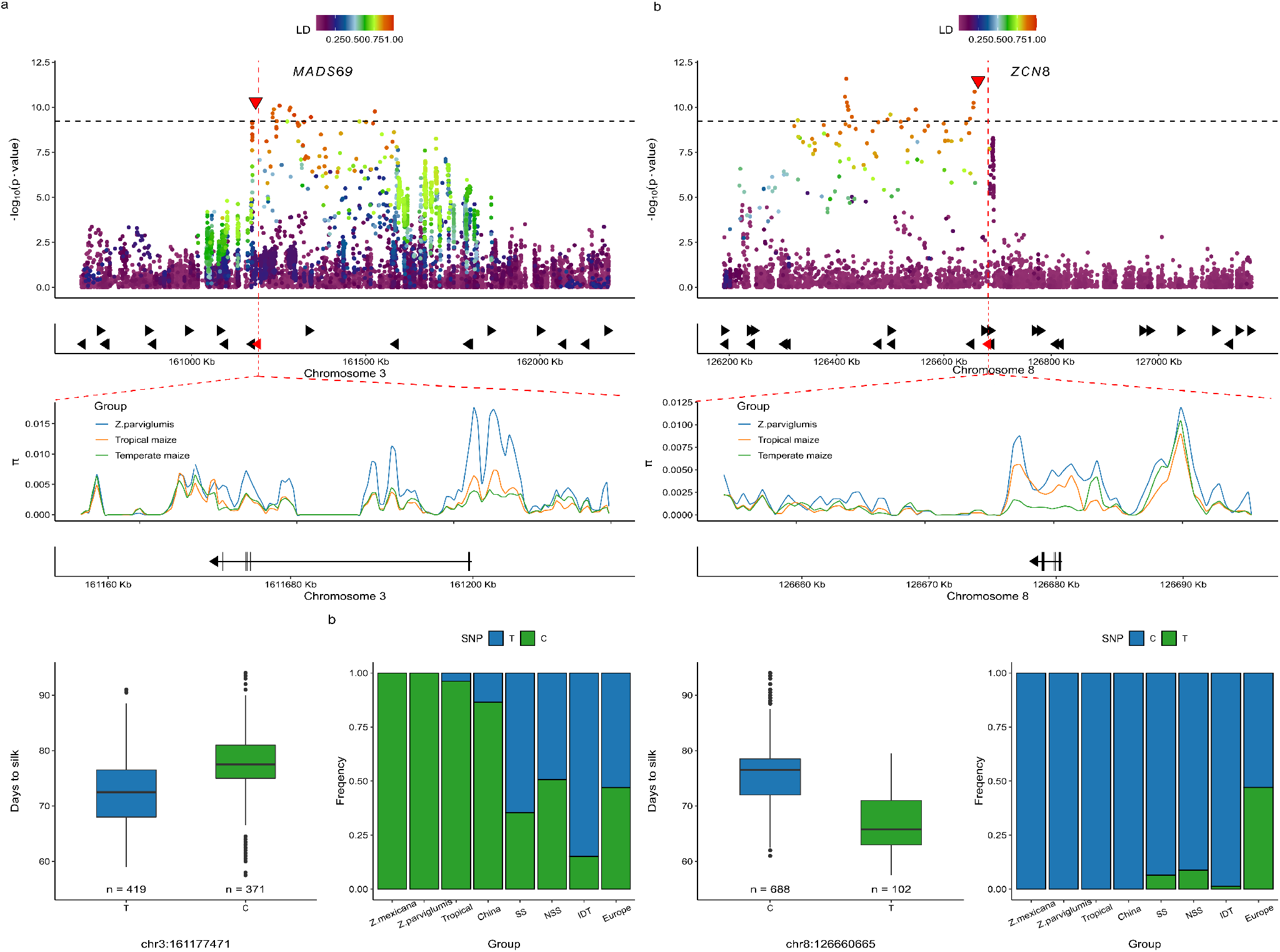
*MADS69* and *ZNC8* are associated with flowering time and were target of selection. Top panel: zoom in on GWAS peak around *MADS69* (**a**) and *ZNC8* (**b**). Linkage disequilibrium (LD) were calculated in each loci against top associated SNP: chr3:161,177,471 and chr8:126,660,665 (marked as red triangles). Horizontal dashed line indicates genome wide Bonferroni correction level. Vertical red dashed lines mark the position of the gene of interest. Middle panel: nucleotide diversity in three maize groups. Gene body of *MADS69 ZNC8* were marked at the bottom. Bottom left panels: Allele effect of chr3:161,177,471 (**a**) and chr8:126,660,665 (**b**) on DTS. Bottom right panels: Changes of allele frequency of chr3:161,177,471 (**a**) and chr8:126,660,665 (**b**) in eight maize groups. SS - stiff stalk, NSS - non-stif-stalk, IDT - iodent.

The diverse composition of the population used for genotyping in this study creates an opportunity to detect patterns of selection in the genome and track changes in favorable allele frequency of variants associated with traits of interest, during domestication, adaptation to a new environment, or genetic improvement during modern breeding. Since flowering plays an important role in local adaptation, we attempted to evaluate patterns of selection around the two known flowering time genes identified above. We observed a clear reduction in nucleotide diversity in the promoter of *MADS69* in tropical and temperate maize lines relative to *parviglumis* (Figure 4a), consistent with previous report (Liang *et al*., 2019). The most significantly associated SNP for days to silking in the *MADS69* gene region was located at position 161,177,471. The reference allele (T) was associated with more rapid female silking relative to the alternate allele (C), with a mean difference of ∼5 days (Figure 4a). In both *mexicana* and *parviglumis* populations only the slower flowering C allele was observed (Figure 4a). In lines classified as belonging to the tropical or Chinese populations, the C allele was predominant. In contrast, the T allele was the more common in the three North American populations (stiff stalk, non-stiff stalk, and iodent). The T allele was particularly common among lines classified as belonging to the iodent heterotic group. Similarly, the T allele also made up the majority of genotype calls among European maize lines included in this study. The large increase of frequency of shorter flowering T allele in temperate adapted lines is consistent with strong selection on *MADS69* during maize adaptation to temperate climates.

The second known flowering time gene identified in this study was *ZCN8*, which has been previously shown to contribute to maize adaption to temperate climates and to have experienced a decline in nucleotide diversity in domesticated maize relative to wild teosinte accessions which is in the parity with the conclusion that *ZCN8* was likely a target of selection during maize domestication (Guo *et al*., 2018). However, the greater representation of different maize groups included in this study enabled the more specific identification of a decline in nucleotide diversity specifically between temperate and tropical domesticated maize populations, while tropical maize retained similar diversity to teosinte at this locus (Figure 4b). This result is consistent with selection on *ZNC8* occurring during adaptation to temperate conditions rather than during domestication.

The single most significant marker at the *ZCN8* locus was a C/T SNP at position 126,660,665 on chromosome 8. The T allele appears to be the derived allele and the median line homozygous for T at this position flowered 10.7 days earlier than the median line homozygous for the C allele. While additional markers at the *ZCN8* locus that were not in LD (<0.2) with the most significant marker also exhibited a statistically significant association with flowering time, a haplotype based model that incorporated information from the top not-in-linkage SNP (chr8:126,689,419) did not significantly improve the predictive ability for flowering time vs a single marker model. The rapid flowering allele at *ZCN8* was observed at extremely low frequencies in North American temperate germplasm. Almost all individuals homozygous for the rapid flowering allele originated in Europe (Figure 4b), demonstrating the importance of sampling broader global germplasm pools to have greater power to identify functional variants primarily segregating in only individual geographic regions.

## Conclusion

In summary, we perform a large-scale joint variant calling for 1,515 maize individuals, which include a wide range of maize accessions from multiple continents and eras and discover more than 46 million high-confidence sequence variants. In addition to releasing new sequence data for 239 new maize inbreds, we also release raw and filtered variant lists as well as processed GenomeDB files that will allow this SNP set to be further extended and expanded without the need to realign previously processed samples to the maize reference genome. We have shown that the new variant set accurately describes the population structure used in this study and improves power in genome wide association studies relative to the previous state-of-the-art marker datasets for a large maize association panel.

## Acknowledgements

This project was supported by U.S. Department of Energy, Grant no. DE-SC0020355, the National Science Foundation under grant OIA-1826781, USDA-NIFA under the AI Institute: for Resilient Agriculture, Award No. 2021-67021-35329, the Foundation for Food and Agriculture Research Award No. 602757, and National Science Center (NCN), Poland, Grant no. 2012/05/B/NZ9/03407 and 2017/27/B/NZ9/00995. This project was completed utilizing the Holland Computing Center of the University of Nebraska, which receives support from the Nebraska Research Initiative.

## Data availability

The additional resequencing data generated as part of this project has been deposited in the European Nucleotide Archive (ENA) under the study accession numbers: PRJEB56265, PRJEB56295, and PRJEB56320. Raw VCF files for all 366 million variants identified in this study, imputed VCF files for the 46 million quality and minor allele frequency filtered variations identified as part of this study and GATK GenomicsDBs files to enable new SNP calling with additional populations have been deposited at CyVerse and are available for download from: https://datacommons.cyverse.org/browse/iplant/home/shared/Grzybowski_MaizeSNPset_2022

Please note that the link above will be augmented with a permanent DOI upon publication.

## Author contributions

JCS and MWG conceived the study. JT conducted experiments and generated data. JY and GX provided advice and feedback on the design of experiments and analyses. MWG and RVM designed and conducted analyses and visualized the results. MWG, RVM and JCS composed the initial draft of the manuscript. All authors contributed to writing and editing and approved the final version of the manuscript.

---

**Table S1. Summary of sequenced lines**. Provided as Excel file.

**Table S2.** Comparison of SNPs yielded in this study with those in Hufford *et al*. (2021)

| Inbred line | Concordance rate |
| --- | --- |
| B73 | 0.99 |
| B97 | 0.98 |
| CML228 | 0.98 |
| CML247 | 0.98 |
| CML277 | 0.98 |
| CML322 | 0.98 |
| CML333 | 0.98 |
| CML52 | 0.96 |
| CML69 | 0.98 |
| HP301 | 0.99 |
| Il14H | 0.98 |
| Ki11 | 0.98 |
| Ki3 | 0.98 |
| Ky21 | 0.98 |
| M162W | 0.98 |
| MO18W | 0.98 |
| MS71 | 0.98 |
| NC350 | 0.98 |
| NC358 | 0.98 |
| Oh43 | 0.99 |
| Oh7B | 0.99 |
| P39 | 0.98 |
| Tx303 | 0.97 |
| Tzi8 | 0.92 |

**Table S3.** Number of accessions resequenced in the corresponding studies.

| Source | # of Accessions | Target populations | Sequencing Depth |
| --- | --- | --- | --- |
| Brandenburg *et al.* (2017) | 58 | American and European landrace | 18X |
| Bukowski *et al.* (2018) | 214 | Various | 2X |
| Chen *et al.* (2022) | 208 | Teosinte | 22X |
| Chia *et al.* (2012) | 16 | Various | 4X |
| Kistler *et al.* (2018) | 48 | South America landrace | XXX |
| Unterseer *et al.* (2014b) | 26 | European | 16X |
| Qiu *et al.* (2021) | 473 | North American temperate | 10-55X |
| This Study | 228 | North American temperate | 22X |
| This Study 2 | 11 | Eastern Europe (Poland) | 35X |
| Wang *et al.* (2017) | 35 | Central America landrace | 29X |
| Wang *et al.* (2020) | 198 | Chinese inbred | 13X |
| Total | 1515 |  |  |

**Figure S1.**
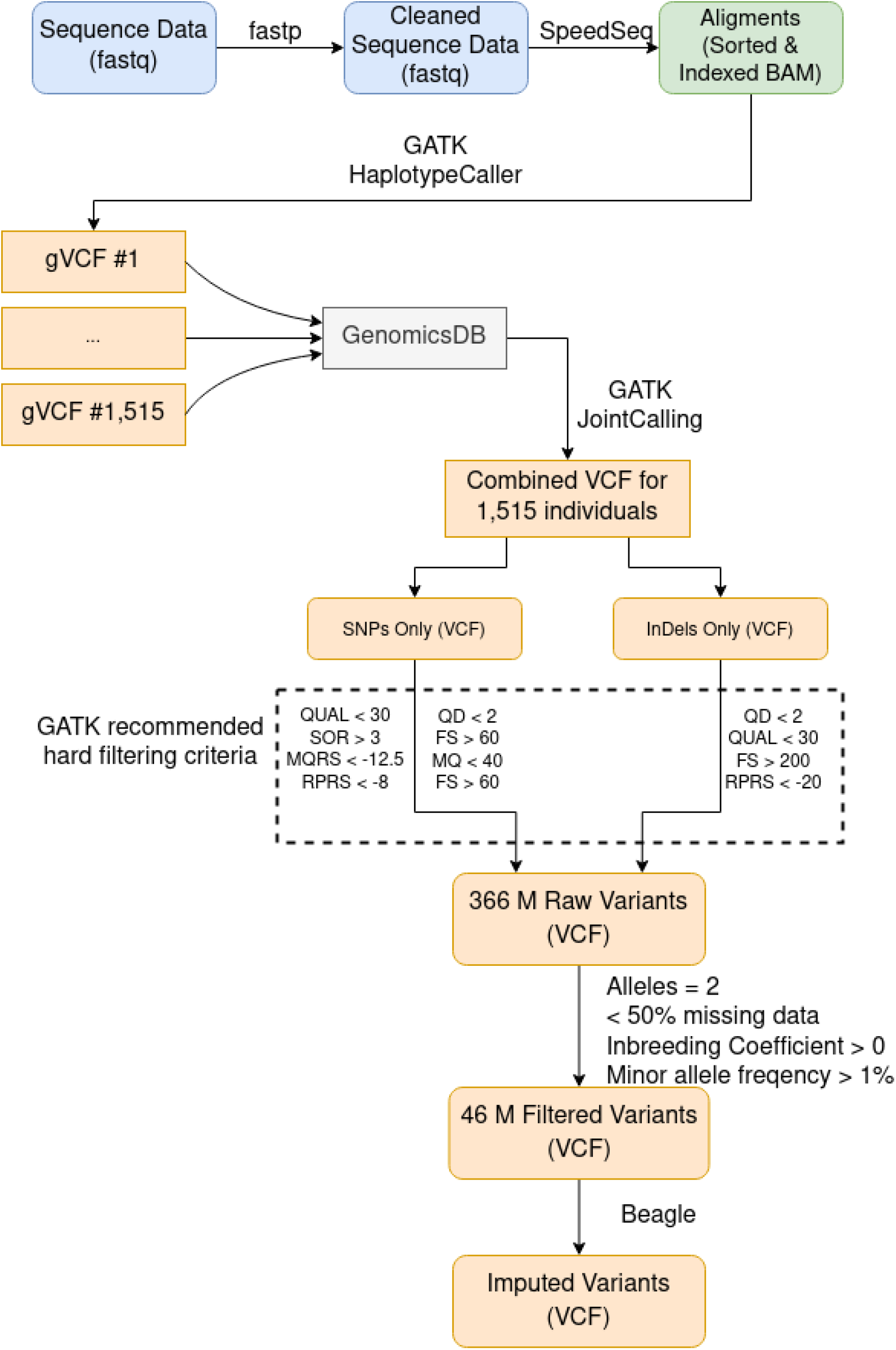
Schematic representation of the approach employed for variant calling and quality filtering in this study.

**Figure S2.**
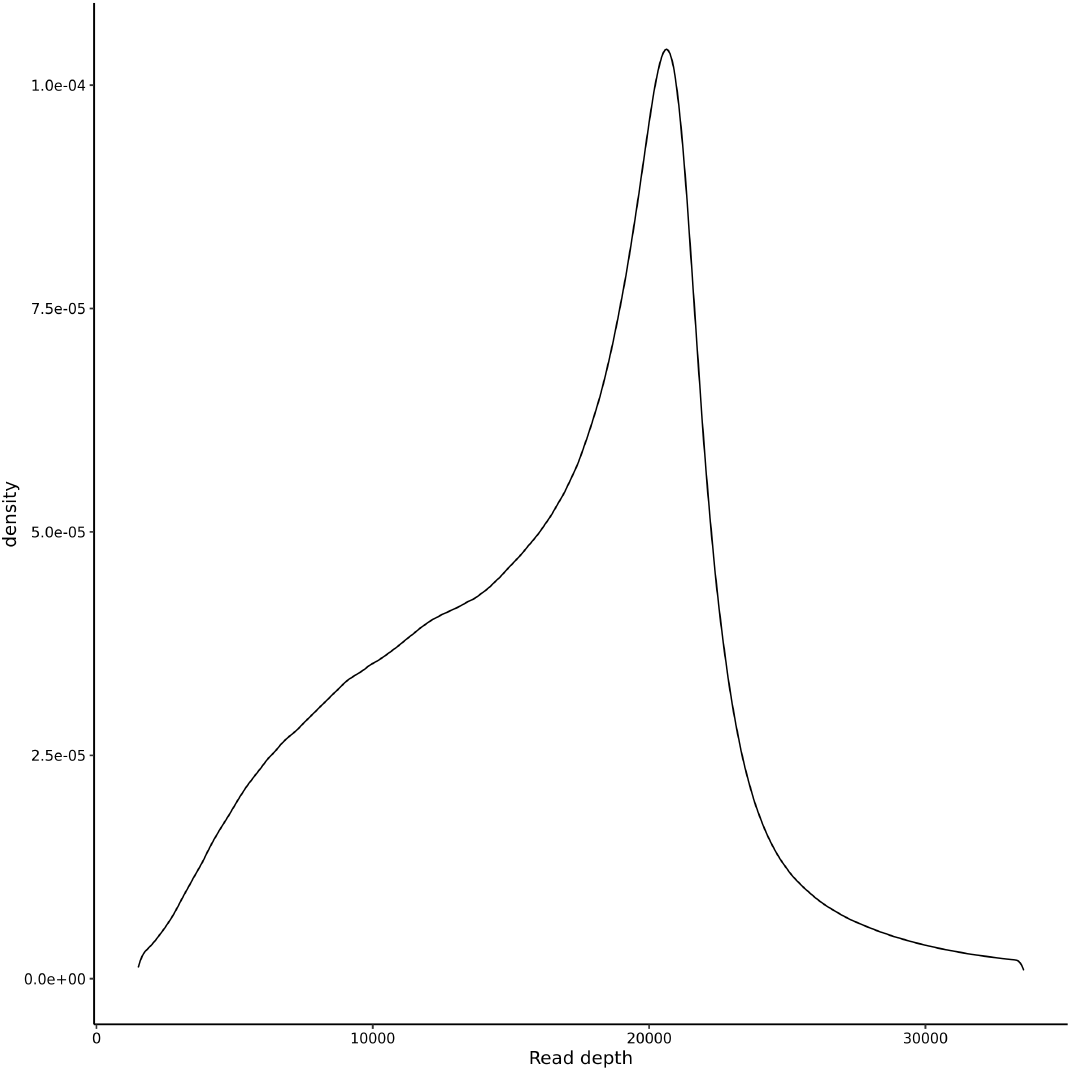
Distribution of total aligned read depth for high confidence (∼46 million) variant sites identified in this study.

**Figure S3.**
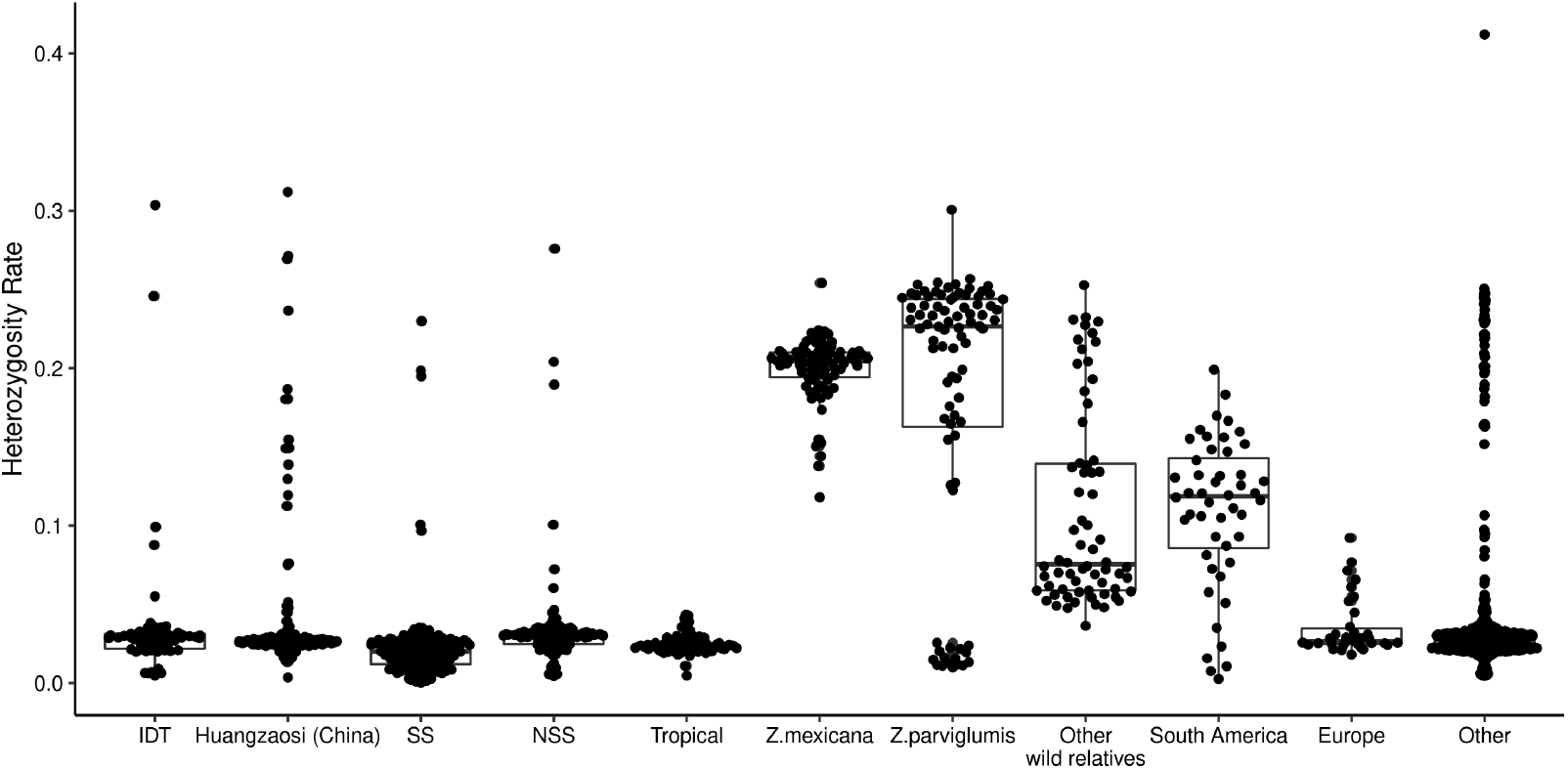
Frequency of heterozygous genotype calls for each individual used in this study. Individuals were assigned to different groups based on previous literature data. Exact heterozygosity rates for each individual plots in this figure are provided in table S1.

**Figure S4.**
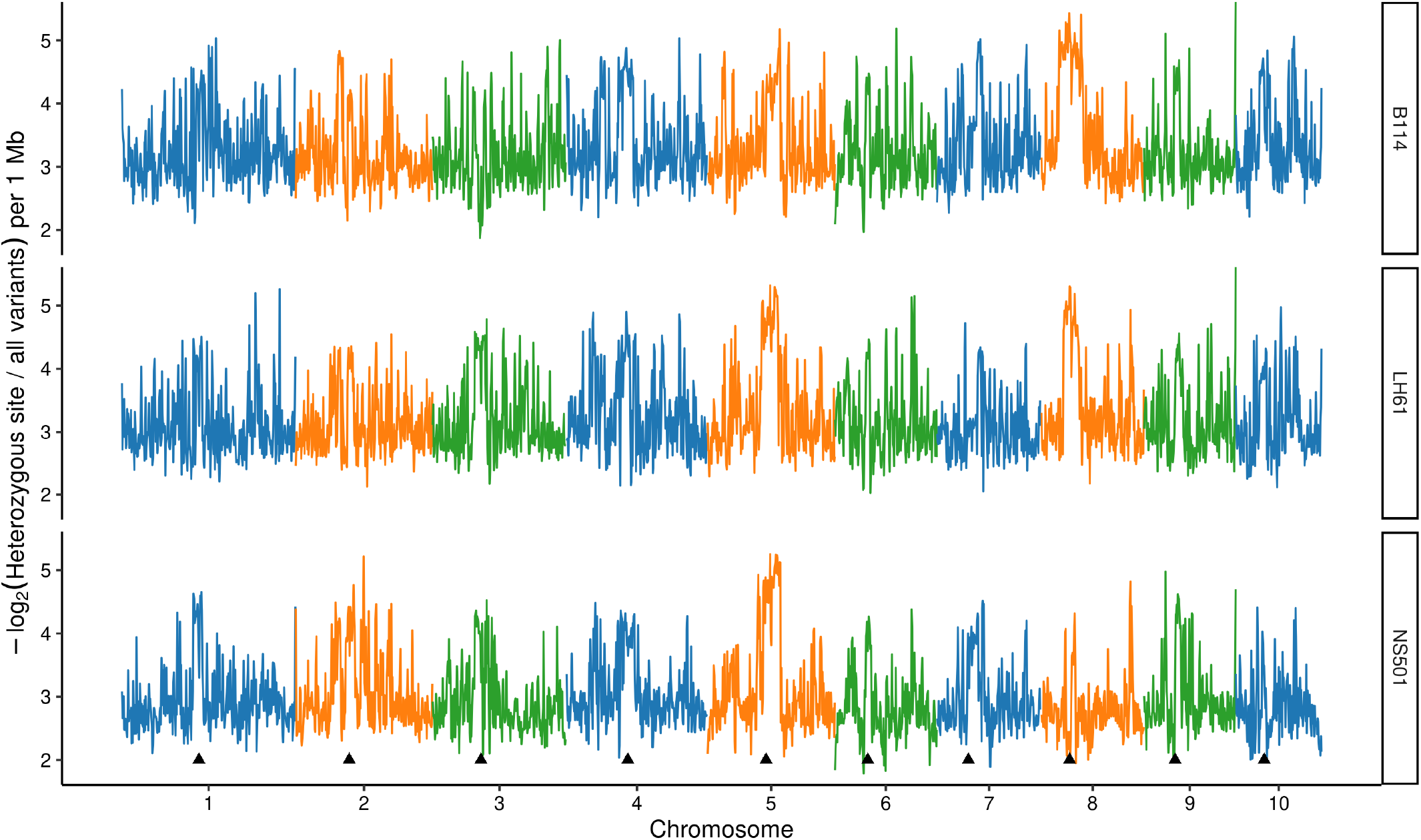
Example of three inbred lines B114, LH61, and NS501 with unexpectedly high heterozygosity. Maize genome was divided into 1 Mb bin and number of heterozygous site in each for each line were counted. Black triangles indicate position of centromere of each chromosome. High confident common (MAF > 5%) variant set were used for this analysis.

**Figure S5.**
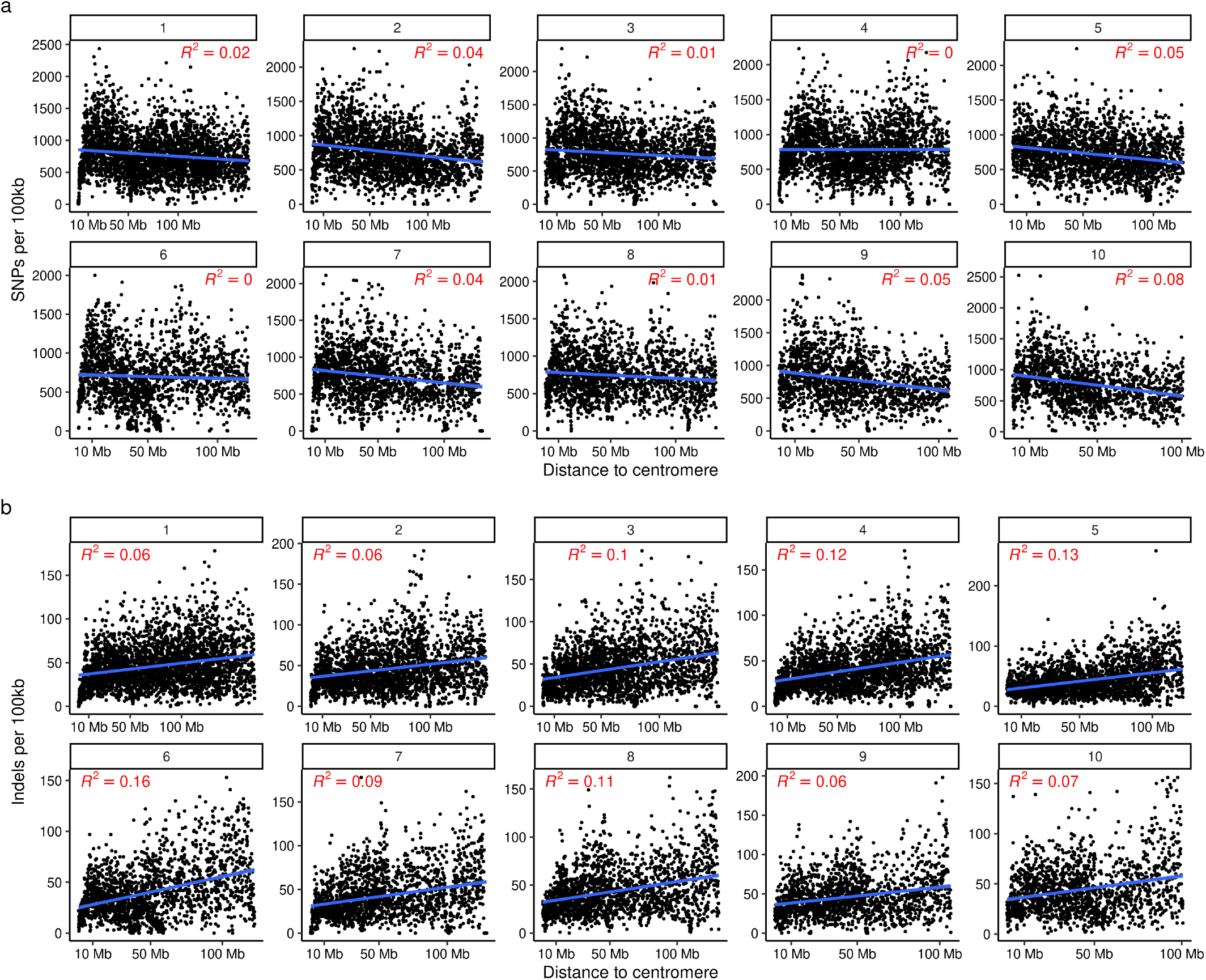
Relationship between distance to centromere and density of SNPs and InDels. Each dot corresponds to a single 100 Kb window on the maize genome, with its position on the x-axis indicating the distance between the bin and the annotated position of the centromere and its position on the y-axis indicating the number of SNPs (a) or InDels (b) present within that interval. Blue line indicates the slope of a linear regression between marker density and distance to the centromere. R^2^ indicates pearson coefficient of determination. All R^2^ values were significant (p < 0.01). The number at the top of each box indicates the maize chromosome number. High confident common (MAF > 5%) variant set were used for this analysis.

**Figure S6.**
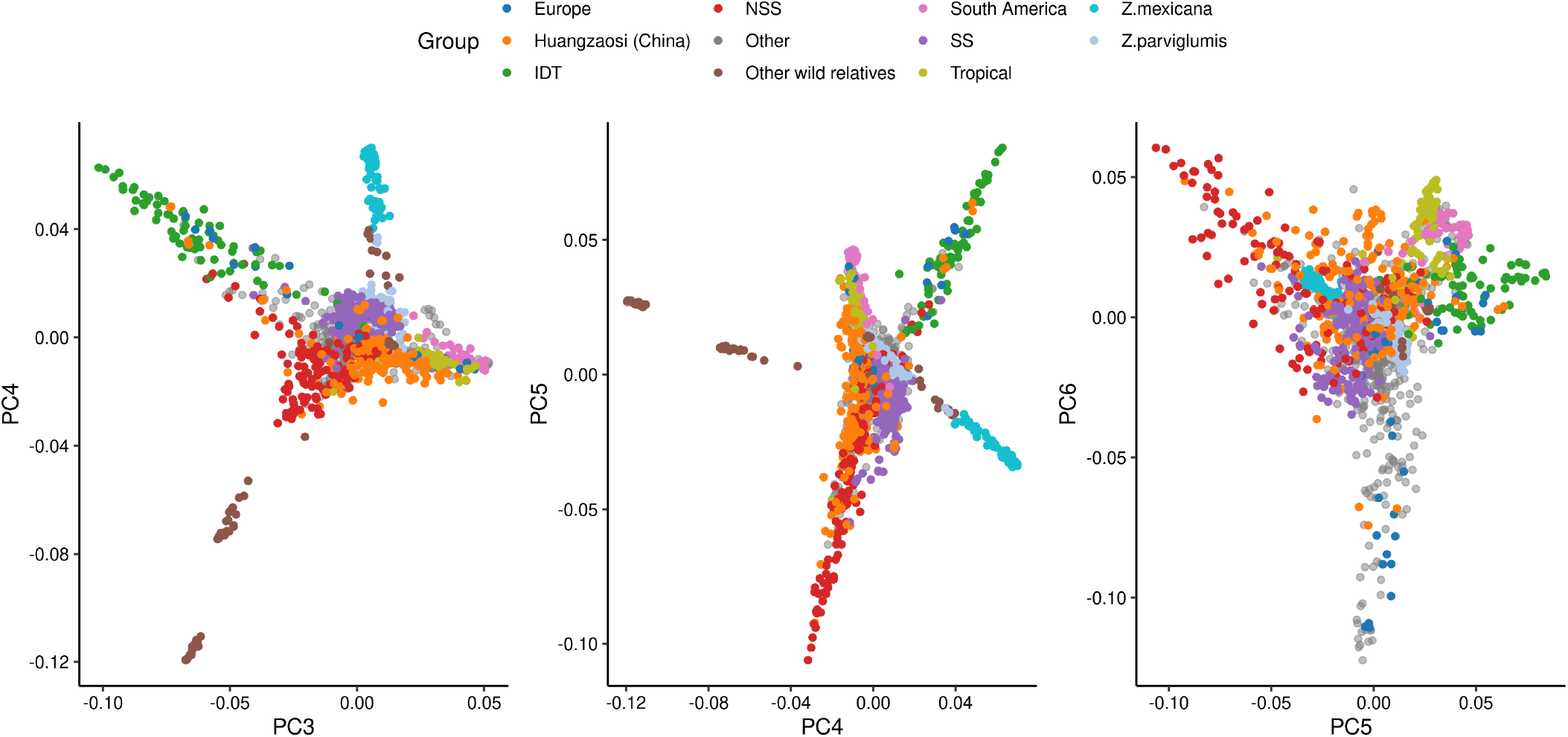
Distribution of PCA values assigned to the 1,515 maize individuals analyzed in this study for principal components three to six. Data are plotted and visualized as described in Figure 2b.

**Figure S7.**
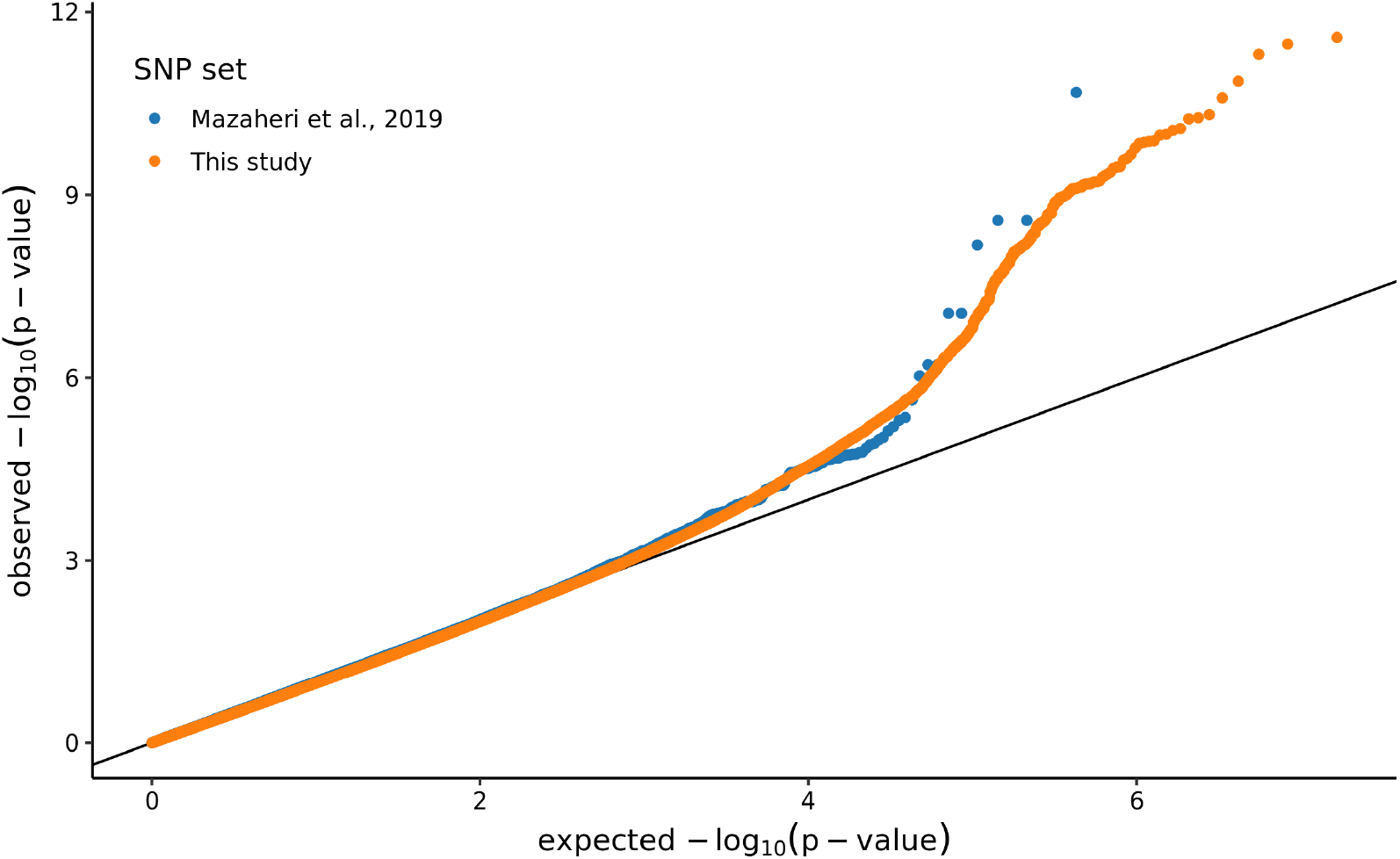
QQ-plots of the two GWAS results shown in Figure 3.

